# Species interactions drive continuous assembly of freshwater communities in stochastic environments

**DOI:** 10.1101/2023.12.21.572946

**Authors:** Andrea Tabi, Tadeu Siqueira, Jonathan D. Tonkin

**Affiliations:** School of Biological Sciences, University of Canterbury, Christchurch, New Zealand; Te Pūnaha Matatini, Centre of Research Excellence in Complex Systems, Auckland, New Zealand

**Keywords:** biodiversity maintenance, causal inference, body size scaling, stochastic modeling, community assembly

## Abstract

Understanding the factors driving the maintenance of long-term biodiversity in changing environments is essential for improving restoration and sustainability strategies in the face of global environmental change. Biodiversity is shaped by both niche and stochastic processes, however the strength of deterministic processes in unpredictable environmental regimes is highly debated. Since communities continuously change over time and space — species persist, disappear or (re)appear — understanding the drivers of species gains and losses from communities should inform us about whether niche or stochastic processes dominate community dynamics. Applying a nonparametric causal discovery approach to a 30-year time series containing annual abundances of benthic invertebrates across 66 locations in New Zealand rivers, we found a strong asynchronous causal relationship between species gains and losses directly driven by predation indicating that niche processes dominate community dynamics. Despite the unpredictable nature of these system, environmental noise was only indirectly related to species gains and losses through altering life history trait distribution. Using a stochastic birth-death framework, we demonstrate that the negative relationship between species gains and losses can not emerge without strong niche processes. Our results showed that even in systems that are dominated by unpredictable environmental variability, species interactions drive continuous community assembly.

## Introduction

Understanding the mechanisms underlying the long-term maintenance of biodiversity is essential for improving conservation efforts and preventing further biodiversity loss (1, 2). Ecological communities change over time and through space as a function of numerous internal and external processes (3). In this continuous assembly process some species persist while others disappear (species losses) and (re)appear (species gains) over time (4, 5) while maintaining local diversity. Various internal and external factors shape community assembly, therefore observing compositional changes over time might shed light on what factors drive assembly processes as well as how biodiversity is maintained in the face of environmental change (5).

The combination of two major mechanisms can lead to continuous community assembly; dispersal-assembly, whereby stochastic processes such as dispersal, random birth and death events dominate (6), and selection- or niche-based assembly (7), whereby species interactions drive community assembly. Ecological communities are on the spectrum between a niche-based and dispersal-based regimes, where their relative positions according to analytic arguments depend on population sizes and the variability of the environment (8). Under highly stochastic environmental conditions, such as in river ecosystems driven by cycles of flood and drought disturbances, community assembly is often assumed to be dominated by external factors such as hydrologic variability that override biotic control of communities (9, 10). However, empirical evidence suggests that species interactions in stream communities, such as competition, predation and herbivory, exert important effects on population and community dynamics (11–13). Species traits such as average body size, voltinism and feeding habits (e.g. predation) have been extensively shown to influence the dynamics of benthic communities (14). In particular, benthic predators often influence the evolution of prey trait distributions. Predatory effects include increase in prey body size and change in body shape, increase in movement speed (15, 16), or change in voltinism such that prey species grow more slowly in the presence of predators (17).

In order to empirically detect the driving force of ecological communities, we investigate the temporal relationship between species (re)appearances (gains) and disappearances (losses) in the community. Community assembly driven primarily by stochastic processes should render species gains and losses independent, uncorrelated events. However, it is possible that the same external factors drive local population disappearances or (re)appearances, potentially leading to correlated gain-loss processes — also known as the confounding effect (18). By contrast, niche-based assembly theory asserts that species diversity arises from ecological selection (i.e. partially or non-overlapping niches). Species fitness differences and interactions are the main drivers of assembly that hypothetically might lead to not only correlated but causally-related species gain-loss events (19). To test this cause-effect relationship, we use time-series data of benthic invertebrate communities from 66 locations across New Zealand recorded between 1990 and 2019 (Fig 1A). River flow regimes are a dominant external force in regulating stream biodiversity (20–22). Therefore, we couple these community data with continuously monitored river flow data and species traits to generate causal linkages. Due to its maritime climate, New Zealand running waters are highly unpredictable and aseasonal relative to continental systems (23, 24). First, we establish our causal hypothesis for how species gains and losses are related and how external processes, in the form of highly dynamic river flow regimes, regulate this connection. To discover these causal hypotheses, we employed a nonparametric causal discovery approach based on conditional independence testing (18, 25). Since observational data are often confounded, they fail to establish cause-effect relationships. In this line, causal inference tools have been developed that allow us to infer causation from observational data (18, 25). Then, based on the discovered causal links we use a stochastic dynamic model combined with scaling theory (26) to theoretically investigate what processes can potentially generate the observed cause-effect relationship between species gains and losses.

**Figure 1.**
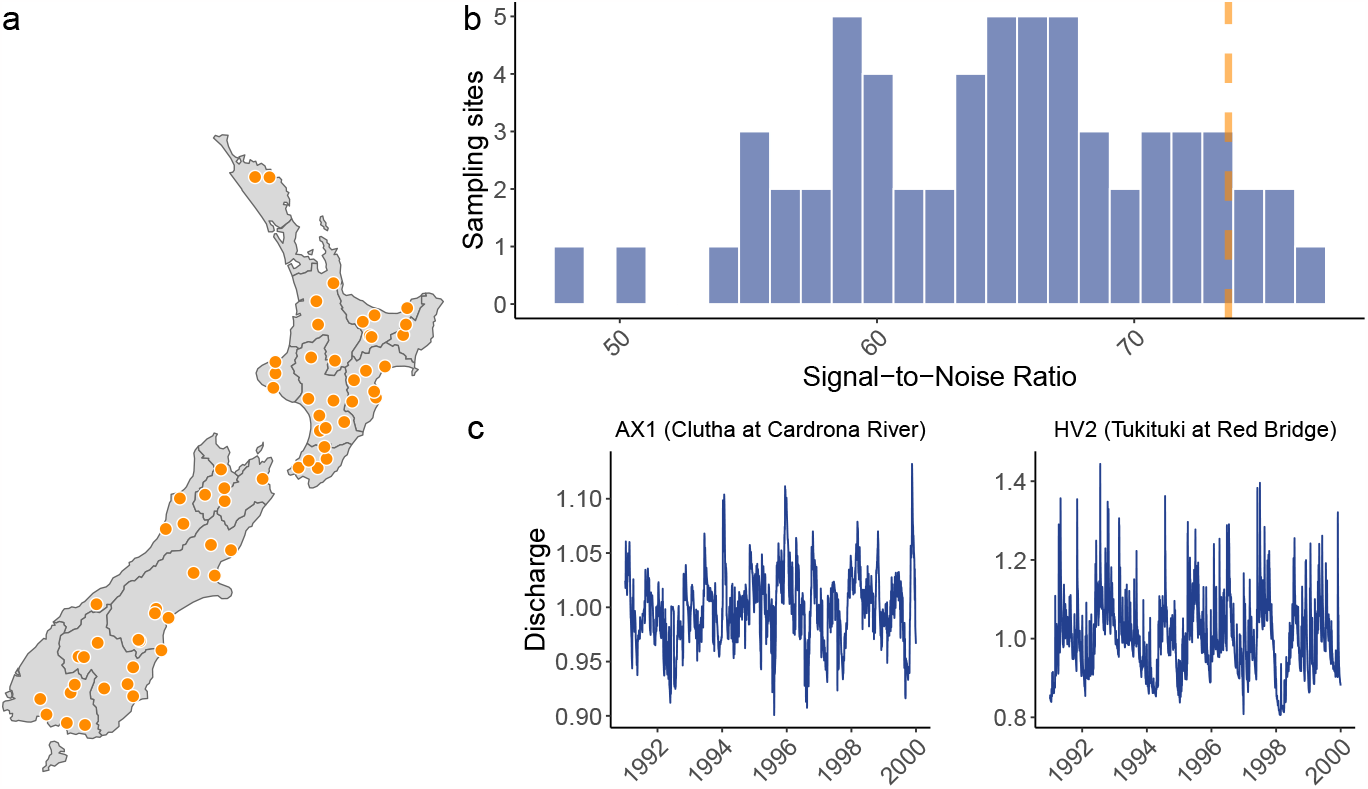
Sampling locations and river discharge in New Zealand. **(a)** Macroinvertebrate communities sampled annually across New Zealand rivers over 30 years at 66 sampling sites. **(b)** The signal-to-noise ratio (SNR) of river discharge time series indicate that New Zealand have rivers ranging from **(c)** aseasonal (left) to highly seasonal (right) discharge patterns. The orange vertical dashed line indicates the SNR of the highly seasonal Yampa river (Colorado, US) for comparison.

## Results

### Empirical results and causal hypothesis

Despite the highly unpredictable river flow regimes observed in these systems, we found clear evidence of deterministic forces structuring the benthic macroinvertebrate communities. The signal-to-noise ratio of river flow indicates that macroinvertebrate communities experience a highly stochastic environment in most rivers (Fig 1B-C). However, benthic communities showed relatively stable biodiversity patterns over time (Fig 2A-C) — we found that relative species richness increased only slightly, while species evenness and species turnover remained constant during the entire time period.

**Figure 2.**
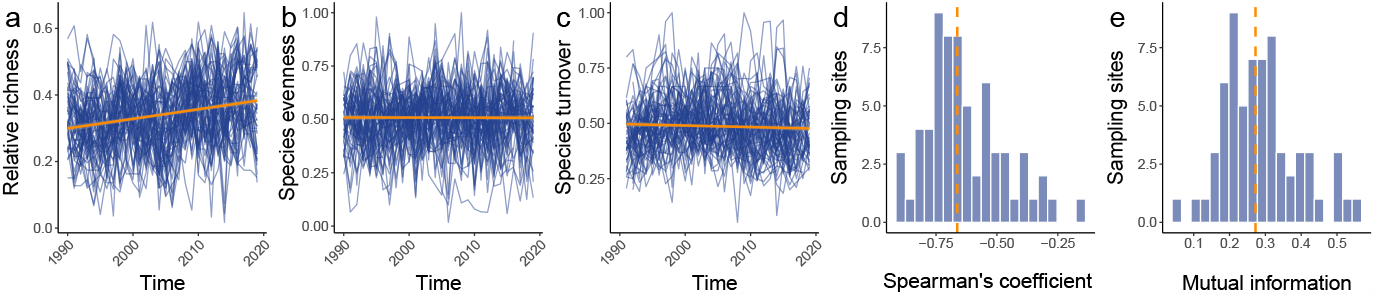
Community metrics. Macroinvertebrate communities were sampled annually across New Zealand rivers over 30 years at 66 sampling sites. **(a)** Communities show overall a slight increase in richness through time. **(b)** Species evenness was also unchanged through time. **(c)** Species identity changes in communities were steady over time with relatively high turnover rates. **(d)** Species gains and losses were negatively correlated in each community (measured as the Spearman’s correlation) textbf(e) with various levels of mutual dependence.

Species gains and losses — defined as the relative number of species gained and lost from one time period to the next (27, 28) — showed strong negative associations and high mutual dependence in all locations (Fig 2D-E). However, to gain cause-effect knowledge about the relationship between species gains and losses that could potentially point towards the importance of species interactions, a context must first be established. Specifically, we assume that this context incorporates stochastic processes such as environmental noise and dispersal along with other species traits that potentially affect species gains and losses in relation to assembly processes. Because species traits are naturally not independent of each other, but rather inter-related, graphical models are needed to establish causal relationships and accurately estimate effect sizes. This graphical model — also called as a causal hypothesis — can be constructed using expert knowledge or intuition or by means of causal discovery algorithms (18). Here we applied a causal discovery algorithm (29) on concatenated time series of each variable from all sampling sites in order to obtain a reliable average estimate of each causal link. Environmental noise was calculated from river flow data (30) and species traits were measured as the community weighted mean (CWM) of body size, voltinism, dispersal, and predation (for details see Methods). The direct structural causal effects between two variables were quantified as partial Spearman’s correlation coefficients (18). The causal analysis indicated that species gains and losses are causally related, describing an asynchronous behavior (Fig 3), which was partly driven by predation. The removal of predation from the causal analysis disconnects gains and losses from the rest of the graph, which indicates that predation is the only community trait directly connected to gain-loss cycles. Surprisingly, environmental noise from river flow was not directly related to species gains and losses, but slightly increased the average generation time in the community. Body size was strongly connected to all other species traits with similar effect sizes confirming its importance of body size in structuring communities. As expected, predation and dispersal affected body size distributions in opposite directions; predation led to larger average body sizes, while better dispersal abilities caused smaller average body sizes in the community. In turn, larger-bodied communities resulted in longer average generation time.

**Figure 3.**
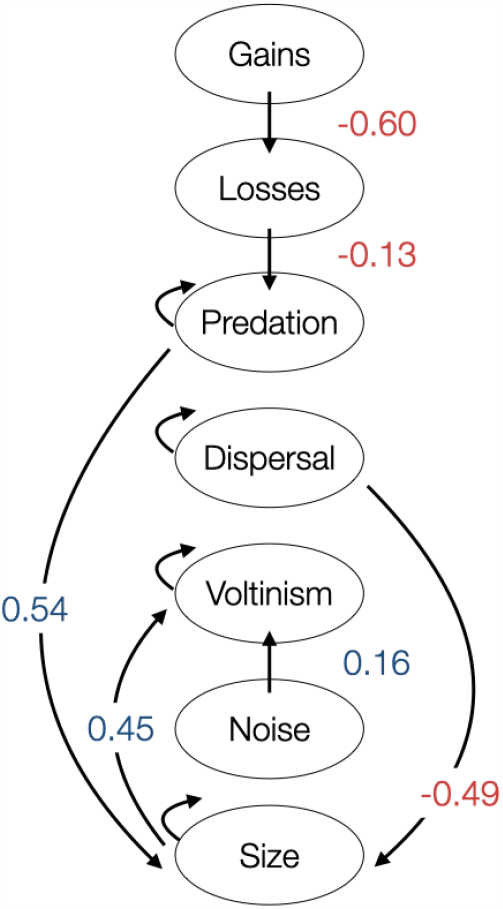
Causal graph. Causal relationships between species gains and losses, environmental noise (measured as the noise component of Fourier transform of river discharge) and community weighted mean (CWM) traits. Using causal discovery (PC algorithm) for time series data, results show that predation is the only variable that is directly linked to species gains and losses. Species gains and losses have a negative bidirectional relationship indicating the presence of cycles. Higher environmental noise slightly reduces predation and also increases the mean generation time (voltinism) in the community. Higher predation leads to larger average body sizes and higher dispersal tend to lead to smaller body sizes. Larger average body sizes increases the average generation time.

### Theoretical results

Based on the findings of the causal inference analysis, we can now establish a simple theoretical investigation in order to determine how the observed patterns between species gains and losses were generated given the relative strength of biotic interactions and stochastic processes. We generated synthetic communities combining a stochastic dynamic model with metabolic scaling theory (26, 31). Specifically, we defined stochastic population dynamics as a birth-death process, where new individuals are gained by *B*(*n*_*i*_) = *q*_*i*_ + *n*_*i*_ *λ*_*i*_ ·(1 − *n*_*i*_*/K*_*i*_), where *q*_*i*_ is density-independent immigration rate and *λ*_*i*_ is the intrinsic growth rate. Species lose individuals as *D*(*n*_*i*_) = *d*_*i*_ *n*_*i*_ + *n*_*i*_ Σ(*a*_*ij*_ *n*_*j*_*/K*_*i*_), where *d*_*i*_ is death rate, *K*_*i*_ is the carrying capacity and *a*_*ij*_ is the interaction coefficient including competitive and predator-prey interactions (see Methods for details). First, we randomly generated interaction matrices, where predation and competition coefficients were determined based on body size scaling (32, 33). Second, dispersal processes were mimicked by the immigration rate (*q*_*i*_), which was set according to the discovered relationship between body size and dispersal in the causal analysis. The intrinsic birth rates (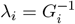, *G* generation time) were also set according to the discovered relationship between size and voltinism (see Methods and Fig S5). Then, each community was sampled over 30 times by equal intervals under different levels of average interaction strengths (*μ* = {0, 0.1, 0.5, 1}) defined as the mean value of all interspecific interaction coefficients. Note that interaction matrices contain both positive and negative coefficients and pairwise interactions are asymmetric. The carrying capacity (*K*_*i*_) was also scaled with body size and we assumed the same death rate (*d*_*i*_) for each species stemming from external sources such as flooding. All community started from a regional species pool with 55 species that is median value of empirical observations (Fig S4). The theoretical analysis confirmed that the observed empirical patterns are driven by strong species interactions. When the average interaction strengths are higher, the relationship between species gain and losses are stronger compared to very low level of interspecific interactions (Fig 4A-B). Relative richness, evenness and turnover approached the observed values when interactions were stronger (Fig 4C-D). Stochasticity alone, i.e. where the average interactions strength is zero, leads to the correlation between species gains and losses approaching zero as well as to high relative richness, highly-even species distributions with low species turnover. As expected, imposing stronger interactions reduces local species richness and species evenness and increases species turnover moving all metrics closer to the observed values.

**Figure 4.**
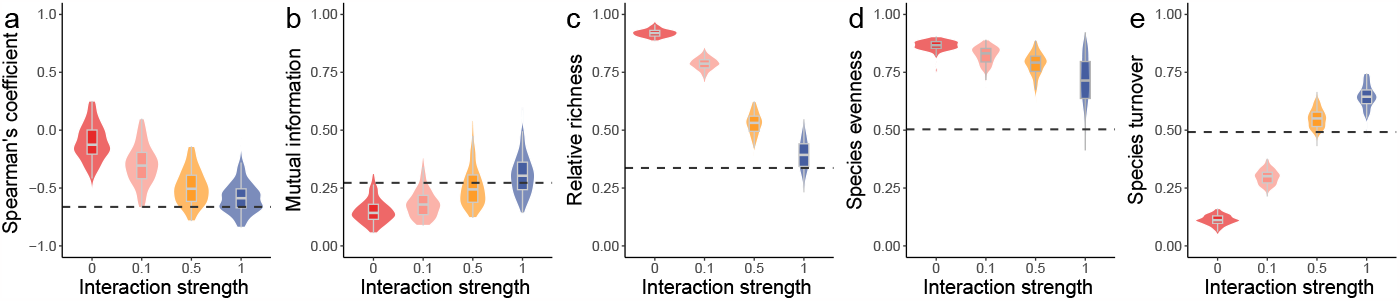
Synthetic analysis of continuous community assembly. Communities assuming stochastic birth-death processes were generated with different levels of interaction strengths over 30 sampling events repeated 100 times for each parameter level. Interaction strengths refer to the average value of the non-diagonal elements of the interaction matrix. The purple dashed line indicates the empirical values. Panel **(a)** shows the Spearman’s correlation coefficient and panel **(b)** depicts the mutual information between species gains and losses time series. Stronger species interactions caused a more stronger negative association between gains and losses time series and higher mutual information. **(c)** Species richness is the highest when species do not interact and decreased with interaction strength. Similarly, interaction strength decreased **(d)** species evenness and increased **(e)** species turnover.

## Discussion

Both empirical and theoretical findings suggest that species gains and losses are causally-related driven by strong biotic interactions in these stochastic environments. Species gains and losses empirically showed asynchronous behavior, similarly to previous observations (28, 34). The asynchronous relationship between species gains and losses is the product of the combination of interaction structure and stochastic processes, whereby a small fraction of species persisted over time, another fraction of species had an intermediate temporal presence and the remainder species rarely appeared potentially resulting from stochastic processes and weak competitive abilities. Our theoretical predictions based on body size scaling relationships also supported that biotic interactions are needed to reconstruct the observed asynchronous relationship between species gains and losses. The role of biotic interactions in the dynamics of river ecosystems have been long debated because of the strong external forcing from cycles of floods and droughts (9, 10) and are therefore deemed to be highly-context dependent (14). For instance, while previous work found that flow variability breaks down competitive hierarchies (11) and predator-prey interactions (35), benthic predators have been suggested to have cascading effects on altering prey abundance, size or age structure, behavior, and morphology (14). Our causal inference analysis identified predatory effects to be partly responsible for the observed continuous community assembly reflected by species gains and losses. Second, the synthetic analysis strongly supported the role of predator-prey and competitive interactions shaping community dynamics closely matching the empirical observations.

We showed that environmental stochasticity affects the number of generations per year (voltinism),i.e. communities under higher environmental noise comprised more species with longer generation time. However, more precise information on changes in voltinism in low and high environmental noise requires further investigation with measured species trait distributions. We observed a limited effect of environmental noise on the communities in these dynamic rivers. This weak influence is expected in living systems due to species adapting to the fluctuation structure of their environment by observing and learning its statistics (e.g. variances and correlations) given that it remains constant over evolutionary timescales (36). In our case, the highly autocorrelated noise with relatively small or in some cases nonexistent characteristic signal present in stream flow measurements (Fig 1B) suggests that species will have developed adaptive strategies such as bet hedging (37). For instance, most predatory species in our analysis were also generalists suggesting an adaptive feeding behavior to a constantly-changing environment. Due to its maritime climate and unpredictable flow regimes, New Zealand stream communities are a case in point of such adaptation, being highly generalist and opportunistic (23).

Our synthetic analysis generated predictions tightly coupled to the observed metrics. As expected, increasing internal constraints reduced the number of species present from the regional pool and reduced evenness due to stronger predation and competitive exclusion within the communities. The presence of internal structure led some species to persist and some species to disappear and reappear according to stochastic events, which creates the observed asynchronous fluctuations of species gains and losses. When species weakly interact, more species were included in the local communities from the regional species pool leading to highly even species distributions and low species turnover, which rendered species disappearances and (re)appearances independent events confirming previous expectations (19). The synthetic analysis also revealed that the discovered causal relationships among species traits in stochastic model communities can be utilized to closely reproduce observed biodiversity patterns, without directly inferring species interactions coefficients from empirical data. In our theoretical investigation, we assumed that biological rates and interactions vary as a function of species body sizes. Body size, as a master trait, is known to scale with other species traits such as dispersal ability (38), predation (39), and voltinism (40). Here we empirically demonstrated that body size not only correlates with, but is causally-related to other species traits and biological processes in stream communities. Predation and dispersal changed body size distribution in benthic communities corroborating previous observations (15, 38). The increase in average body sizes can be explained by size-selective predation of smaller-bodied prey species. Therefore, we assumed that macroinvertebrate communities are size-structured and likely governed primarily by predator-prey and competitive interactions, however, other interactions types such as facilitation might have an important role in macroinvertebrate communities (41). For instance, aggregation, a form of facilitation, reduces the individual risk of predation and can benefit individuals by recycling each other’s byproducts (42). Nevertheless, the causal association between predators and gains and losses, and the lack of any other direct association, indicates a dominant role of antagonistic interactions in structuring these communities.

In this work, we showed that species occurrence information allow the detection of mechanisms driving community dynamics by combining causal inference analysis with theoretical models. Following a causal discovery approach, we identified causal links between species traits, environmental noise and internal processes. Then, we used the information obtained from causal discovery to calibrate and parameterize a stochastic trait-based dynamical model. Given the high match between the theoretical results and observations, we believe that our work provides a future avenue towards a data-driven general framework to investigate continuous community assembly.

## Methods

### Data

Overall, we analyzed 1795 communities from 66 geographical sites (Fig 1) across New Zealand comprising population abundance data from more than 114 macroinvertebrate taxa sampled from 1990 to 2019. These surveys were conducted for New Zealand’s National River Water Quality Network (NRWQN) (43). Samples were collected following standardized protocols (43) and under baseflow conditions. Seven Surber samples (0.1 m^2^ and 250 *μ*m mesh net) were collected on all sampling occasions during which macroinvertebrates were removed from a 0.1 *m*^2^ area in the sampler down to a depth of ca. 10 cm and from as many substrate types as possible. Individuals were later identified in the laboratory, to the lowest practicable taxonomic level (44). The information on functional traits related to morphology, life-history, dispersal strategies and resource acquisition methods was obtained from the New Zealand freshwater macroinvertebrate trait database prepared by NIWA, which has been explicitly developed for New Zealand’s standardised freshwater macroinvertebrate sampling protocols (45). Functional traits were fuzzy-coded from 0 to 3 (46) and converted to a single value for each taxon using weighted averages. Daily average river discharge data (*l/s*) at each sampling location collected from NIWA database. The time series of environmental noise was obtained as the root mean squared noise, or amplitude of all noncharacteristic frequencies of the daily average time series of each year using FFT (Fast Fourier Transform). Flow data were log10-transformed and normalised by the average discharge across the entire period at each site (30).

Species gains and losses were calculated as the number of species gained and lost from previous to next year divided by the total number of species observed in both years (27, 28). Furthermore, we used traditional biodiversity metrics that capture important structural aspects of communities such as number of species at given time compared to the number of species occurred in that given location over the studied period (relative richness), species abundance distribution (evenness) and species identity change over time (species turnover). Species evenness (*J*) is a description of the distribution of species abundances within a community and is defined as the Shannon information entropy divided by the maximum entropy of relative species abundances: 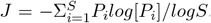, where *P*_*i*_ is the relative abundance of species *i*. Species turnover — defined as the relative number of species gained and lost from one time period to the next — describes local compositional changes over time (27, 28). The relationship between species gain-loss time-series were quantified by Spearman’s correlation coefficient and normalised mutual information. Mutual information is a nonparametric and non-monotonic similarity metric between two random variables and calculated as *MI*_*XY*_ = *H*(*X*) + *H*(*Y*) − *H*(*X, Y*), where *H* denotes the Shannon entropy. The *MI*_*XY*_ was then normalised by the maximum mutual information : 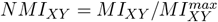, where 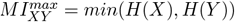

### Causal discovery and causal effects

In order to infer causal relationships from observational time series, a first step involves the construction a graphical causal model, i.e. DAG (directed acyclic graph), among the variables in question referred to as *causal discovery* (18). Causal discovery algorithms based on conditional independence testing (or constraint-based approaches) have four major steps: first, they start with a full undirected graph on *n* nodes (variables), with edges between all nodes. Second, test each pair of variables *X* and *Y*, and each set of other variables *S*, if *X* **⊥** *Y* |*S*; if so, remove the edge between *X* and *Y*. Third, search for *V* -structures (colliders) by checking for conditional dependence and orient the edges of colliders. Lastly, orient the remaining undirected edges (if possible) by consistency with already-oriented edges. We applied the PC (Peter-Clark) algorithm for time series data (29, 47). In order to obtain a general picture of the causal relationships among our variables and to gain statistical confidence, we combined each variable of all 66 sites into a single concatenated time series (48). All variables *V* were shifted with a time lag of *τ* = 1 (variables were first shifted then concatenated). The time lag corresponds to the average generation time of the macroinvertebrate species, where most species are uni- or plurivoltine (Fig S4). We did not include the time lagged version of the environmental noise in the causal model due to its high autocorrelation (Fig S1). Since, all variables were continuous random variables but from different distributions, nonparametric Spearman’s partial correlation test have been applied to test for conditional independence (49, 50). The threshold for conditional independence tests were set to *α* = 5 · 10^−4^ based on SHD (Structural Hamming Distance) analysis (Fig S2) (51, 52). The window causal graph (Fig S3), which covered all variables (*V*_*t*_ and *V*_*t*+1_), showed time consistency. The summary causal graphs (Fig 3) which was deduced from the window causal graph and directly relate variables without time, gives an overview of the relationships (47). The effect sizes were calculated as partial correlation coefficients applying the corresponding adjustment sets based on the window causal graph.

### Theoretical analysis

We defined the population dynamics as a birth-death process (8, 53). The population birth *B*(*n*) and death *D*(*n*) rates are expressed as *B*(*n*_*i*_) = *q*_*i*_ + *n*_*i*_ ·*λ*_*i*_ (1 − *n*_*i*_*/K*_*i*_) and *D*(*n*_*i*_) = *d*_*i*_ *n*_*i*_ + *n*_*i*_ Σ(*a*_*ij*_ *n*_*j*_*/K*_*i*_). We consider that a community of species is characterized by an interaction matrix (*A*), whose elements (*a*_*ij*_) define the direct per-capita effect of a species j on the per-capita growth rate of a species *i*. Note that *a*_*ij*_ and *a*_*ji*_ are not the same. Interaction matrices were generated using scaling relationships based on species body masses: *M*_*i*_ = *M*_0_ 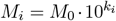 with *k*_*i*_ ∼ *N* (1, 0.3) and *M*_0_ = 1. Competitive and predator-prey interaction coefficients were estimated as 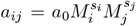 with *s*_*i*_ = 2*/*3 and *s*_*j*_ = 11*/*12 (32) or *s*_*i*_ = −3*/*4 and *s*_*j*_ = 3*/*4 (33), respectively. The immigration rate (*q*_*i*_) scaled with body sizes 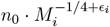 with added Gaussian noise *N* (0, 0.1) to the exponent, where *n*_0_ = 10, which represent the noise resulting from dispersal processes. Note that dispersal abilities is generally positively scales with body mass, however in our empirical analysis smaller macroinvertebrate species have better dispersal abilities. Similarly, the intrinsic birth rates (*λ*_*i*_) were scaled with body masses as 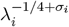 adding noise drawn from normal distribution *N* (0, 0.1) to the exponent. The intrinsic birth rates represent the inverse generation time, i.e. species with longer generation time have lower birth rates. In both cases, the amount of noise added were set to simulate the observed values obtained from causal inference analysis (see Supporting Information). Carrying capacities were calculated as *K*_*i*_ = *K*_0_ 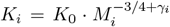, where *γ*_*i*_ is drawn from *N* (0, 0.1) and *K*_0_ = 10^3^. Death rates were uniformly set to 0.5 across all species. In each case, the fraction of predator species were set to 45% similar to observations (see Supporting Information). The state of the system can be characterized by the probability *P* of having *n* individuals at time *t*. The time evolution of the probability distribution is described by a differential equation called a master equation: 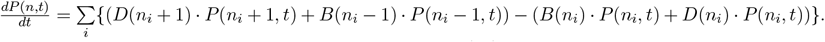 Communities were simulated using Gillespie’s algorithm (54). Simulations were run starting with 55 species representing the regional species pool across different levels of average interaction strengths (*μ* = {0.01, 0.1, 0.5, 1}), each interaction strength was replicated 100 times. Each stochastic simulation process was sampled over 30 times by equal time intervals. At each sampling event species identities and abundances were recorded.

## Acknowledgments

AT was supported by Te Pūnaha Matatini, a Centre of Research Excellence funded by the Tertiary Education Commission, New Zealand. JDT is supported by a Rutherford Discovery Fellowship administered by the Royal Society Te Apārangi (RDF-18-UOC-007).

## Supporting Information

**Figure S1.**
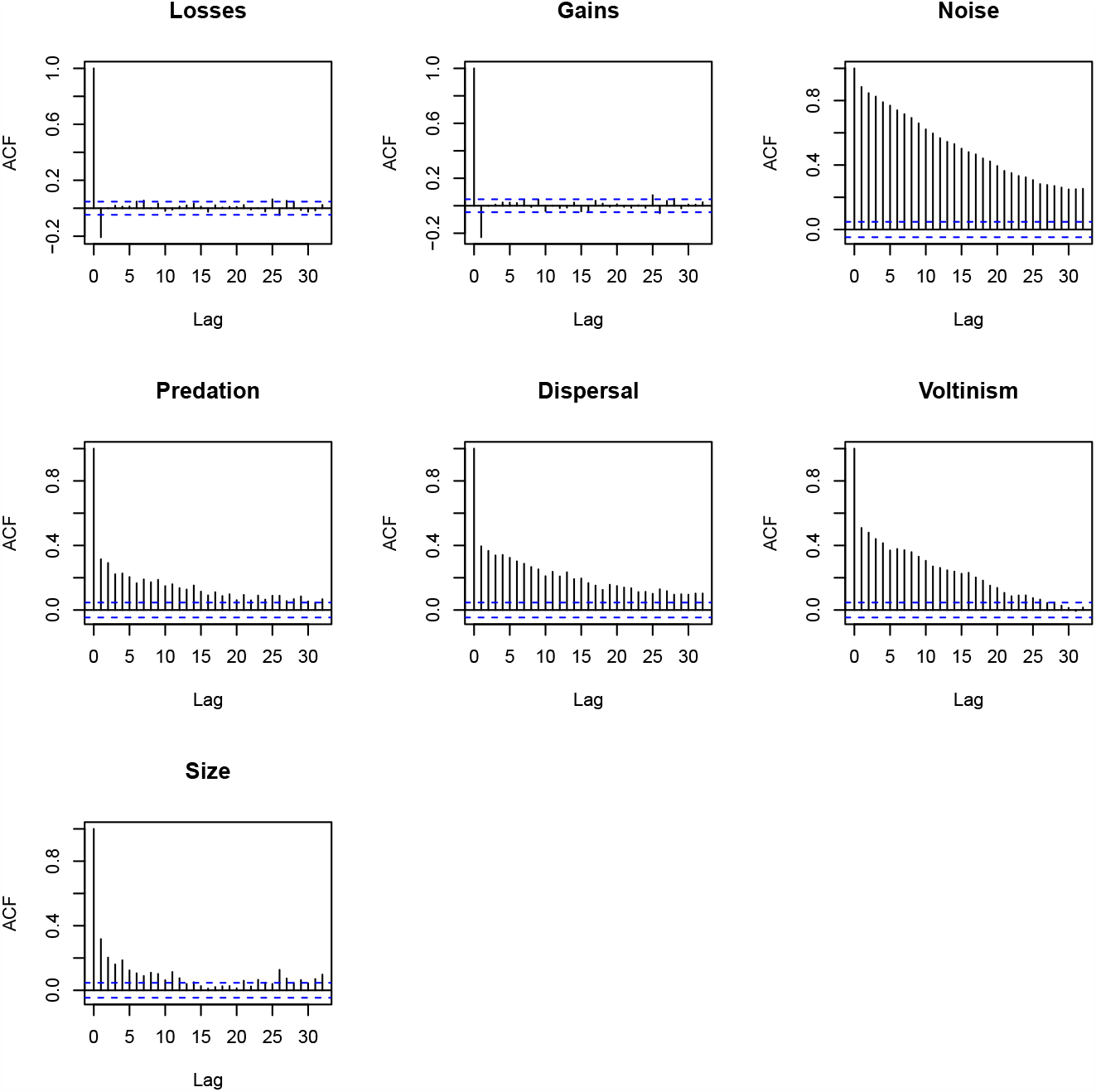
The autocorrelation functions (ACF) for all studied variables. Species gains and losses show negative autocorrelation with 1 year lag. Other variables are positively autocorrelated, especially the environmental noise showing the highest positive autocorrelation.

**Figure S2.**
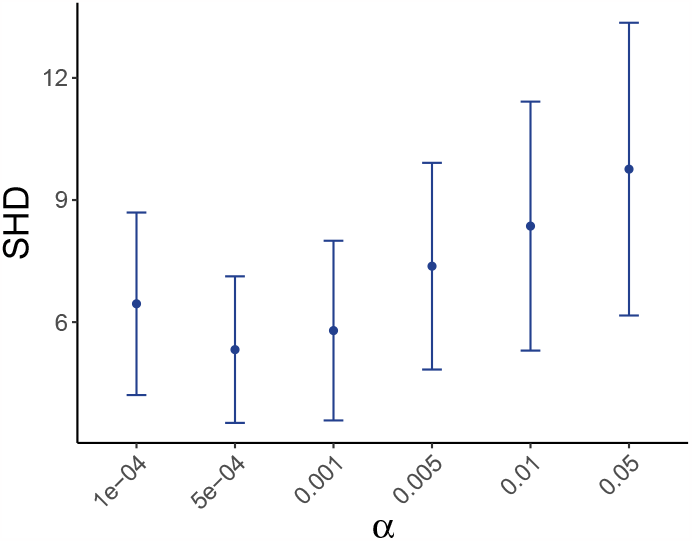
SHD tests across significance levels. Structural Hamming Distance (SHD) were calculated to assess the stability of PDAGs at different significance levels. Based on these results the significance level of *α* = 0.0005 was applied.

**Figure S3.**
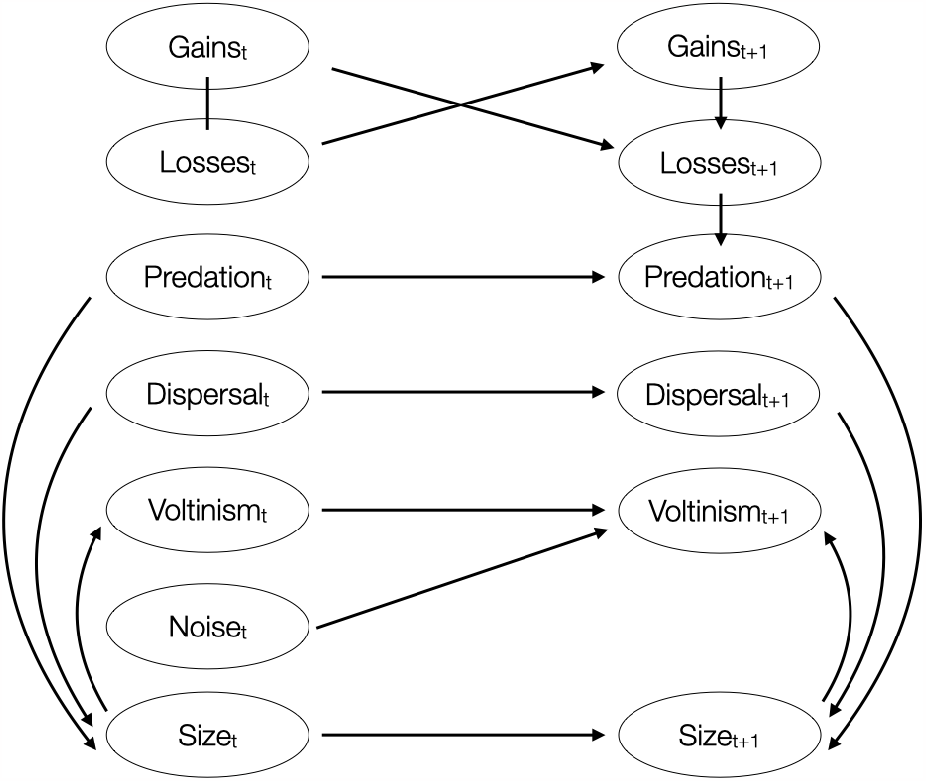
Window causal graph. The full time causal graph shows consistency throughout time. It is a representation of the causal graph through a time window, the size of which equals the maximum lag relating time series in the full time causal graph. All variables and their 1-year lagged version are depicted. The noise variable were not lagged due to its very high autocorrelation.

**Figure S4.**
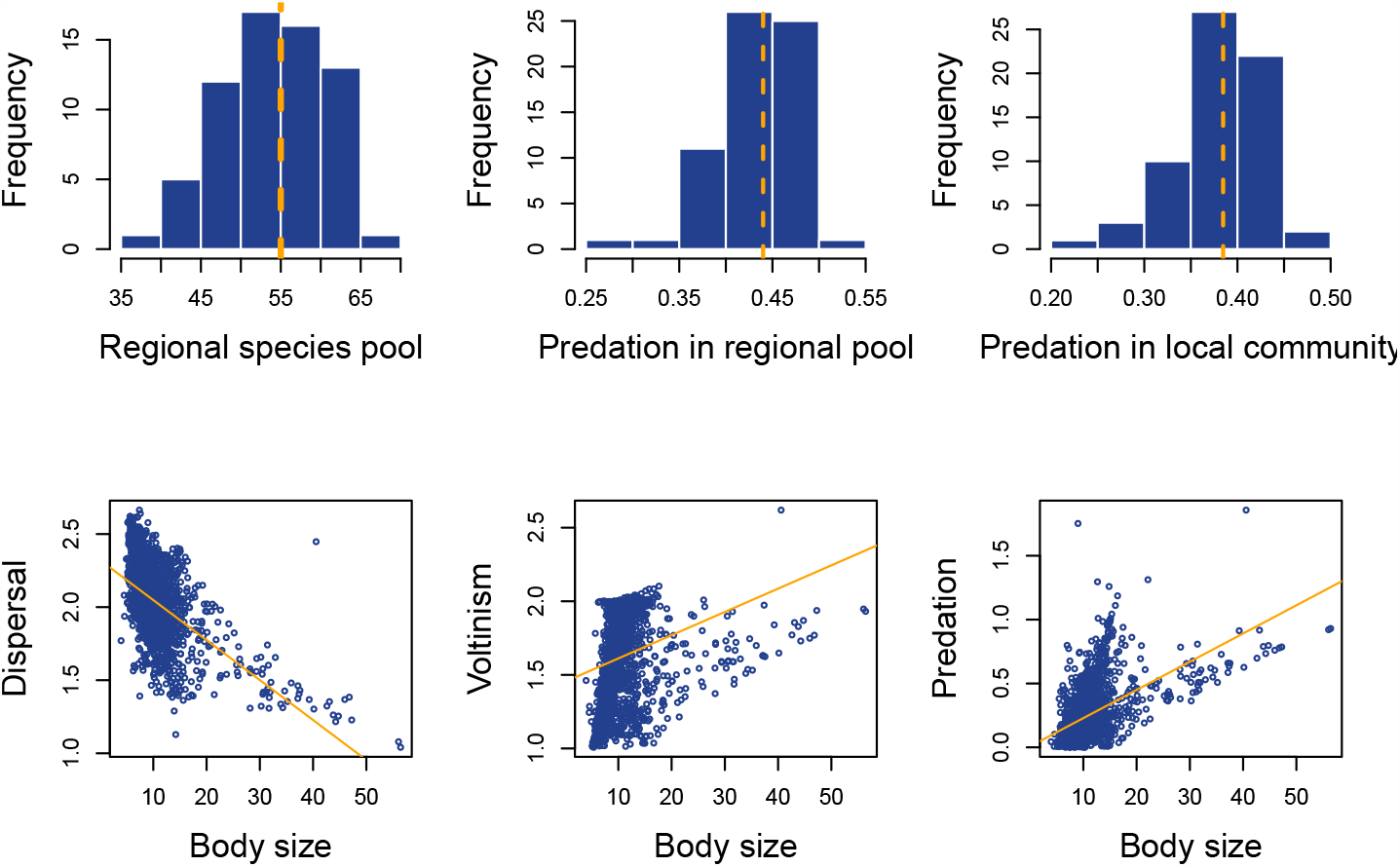
Empirical trait distributions. Regional species pool measured as the number of species appeared over the 30 year period at each sampling site across 66 sites in New Zealand. Relative predation was calculated as the number of predatory species in each regional species pool and as the number of predatory species in each local community. Body size is connected to all studied traits. Voltinism and predation increase and dispersal decreases with body size.

**Figure S5.**
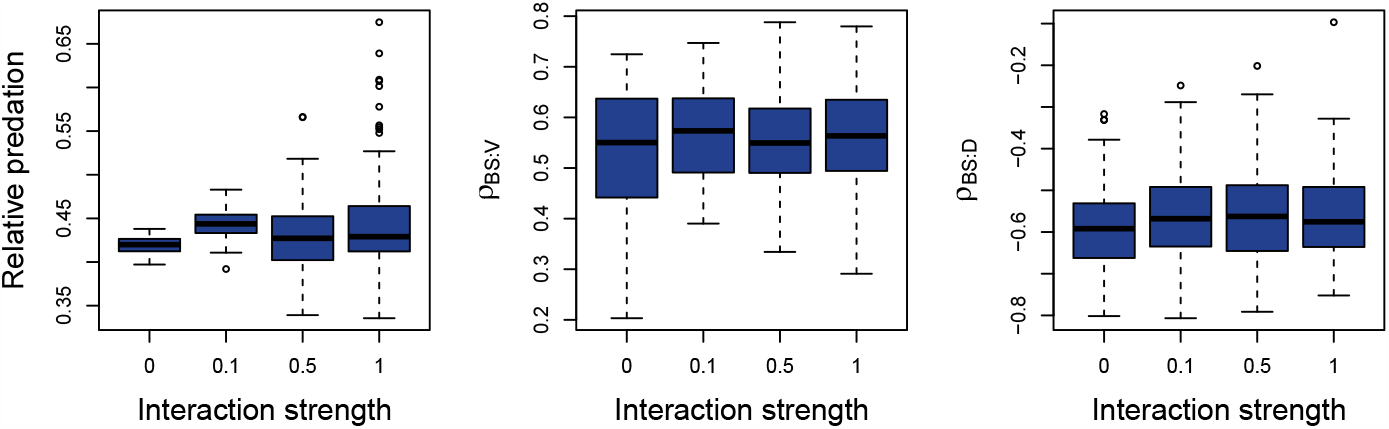
Parameters of simulated distributions. The relationship between body sizes and voltinism (1*/λ*), and dispersal (*q*) were set according to the empirical effect sizes. The number of predators were set to 45% in each simulations and the measured at the sampling events.

## Notes

### Competing Interest Statement

The authors have declared no competing interest.

## References

(1) Sala OE, et al. (2000) Global Biodiversity Scenarios for the Year 2100. Science 287(5459):1770–1774.

(2) Hooper DU, et al. (2005) Effects of Biodiversity on Ecosystem Functioning: A Consensus of Current Knowledge. Ecological Monographs 75(1):3–35.

(3) HilleRisLambers J, Adler P, Harpole W, Levine J, Mayfield M (2012) Rethinking Community Assembly through the Lens of Coexistence Theory. Annual Review of Ecology, Evolution, and Systematics 43(1):227–248.

(4) Nee S, Gregory RD, May RM (1991) Core and Satellite Species: Theory and Artefacts. Oikos 62(1):83–87.

(5) Dornelas M, et al. (2019) A balance of winners and losers in the Anthropocene. Ecology Letters 22(5):847–854.

(6) Hubbell SP (2001) The Unified Neutral Theory of Biodiversity and Biogeography (MPB-32). (Princeton University Press).

(7) Chase JM, Leibold MA (2003) Ecological Niches: Linking Classical and Contemporary Approaches, Interspecific Interactions. (University of Chicago Press, Chicago, IL).

(8) Fisher CK, Mehta P (2014) The transition between the niche and neutral regimes in ecology. Proceedings of the National Academy of Sciences 111(36):13111–13116. Publisher: Proceedings of the National Academy of Sciences.

(9) Poff NL, Ward JV (1989) Implications of Streamflow Variability and Predictability for Lotic Community Structure: A Regional Analysis of Streamflow Patterns. Canadian Journal of Fisheries and Aquatic Sciences 46(10):1805–1818.

(10) Tonkin JD, et al. (2021) Designing flow regimes to support entire river ecosystems. Frontiers in Ecology and the Environment 19(6):326–333.

(11) McAuliffe JR (1984) Competition for Space, Disturbance, and the Structure of a Benthic Stream Community. Ecology 65(3):894–908.

(12) Cooper SD, Walde SJ, Peckarsky BL (1990) Prey Exchange Rates and the Impact of Predators on Prey Populations in Streams. Ecology 71(4):1503–1514.

(13) Rosemond AD, Pringle CM, Ramírez A, Paul MJ (2001) A Test of Top-down and Bottom-up Control in a Detritus-Based Food Web. Ecology 82(8):2279–2293.

(14) Holomuzki JR, Feminella JW, Power ME (2010) Biotic interactions in freshwater benthic habitats. Journal of the North American Benthological Society 29(1):220–244.

(15) Scrimgeour GJ, Culp JM, Wrona FJ (1994) Feeding while Avoiding Predators: Evidence for a Size-Specific Trade-off by a Lotic Mayfly. Journal of the North American Benthological Society 13(3):368–378.

(16) McPeek MA, Schrot AK, Brown JM (1996) Adaptation to Predators in a New Community: Swimming Performance and Predator Avoidance in Damselflies. Ecology 77(2):617–629.

(17) Martin TH, Johnson DM, Moore RD (1991) Fish-Mediated Alternative Life-History Strategies in the Dragonfly Epitheca cynosura. Journal of the North American Benthological Society 10(3):271–279.

(18) Pearl J (2009) Causality: Models, Reasoning and Inference. (Cambridge University Press, USA), 2nd edition.

(19) Spaak JW, Adler PB, Ellner SP (2023) Continuous assembly required: Perpetual species turnover in two-trophic-level ecosystems. Ecosphere 14(7):e4614.

(20) Lytle DA, Poff NL (2004) Adaptation to natural flow regimes. Trends in Ecology & Evolution 19(2):94–100.

(21) Palmer M, Ruhi A (2019) Linkages between flow regime, biota, and ecosystem processes: Implications for river restoration. Science 365(6459):eaaw2087.

(22) Tonkin JD (2022) Climate Change and Extreme Events in Shaping River Ecosystems in Encyclopedia of Inland Waters (Second Edition), eds. Mehner T, Tockner K. (Elsevier, Oxford), pp. 653–664.

(23) Winterbourn MJ, Rounick JS, Cowie B (1981) Are New Zealand stream ecosystems really different? New Zealand Journal of Marine and Freshwater Research 15(3):321–328.

(24) Tonkin JD, Death RG, Muotka T, Astorga A, Lytle DA (2018) Do latitudinal gradients exist in New Zealand stream invertebrate metacommunities? PeerJ 6:e4898.

(25) Shipley B (2016) Cause and Correlation in Biology: A User’s Guide to Path Analysis, Structural Equations and Causal Inference with R. (Cambridge University Press, Cambridge), 2 edition.

(26) Brown JH, Gillooly JF, Allen AP, Savage VM, West GB (2004) Toward a Metabolic Theory of Ecology. Ecology 85(7):1771–1789.

(27) Diamond JM (1969) Avifaunal equilibria and species turnover rates on the channel islands of california. Proceedings of the National Academy of Sciences 64(1):57–63.

(28) Hallett LM, et al. (2016) codyn: An r package of community dynamics metrics. Methods in Ecology and Evolution 7(10):1146–1151.

(29) Spirtes P, Glymour C, Scheines R (2001) Causation, Prediction, and Search. (The MIT Press).

(30) Sabo JL, Post DM (2008) Quantifying Periodic, Stochastic, and Catastrophic Environmental Variation. Ecological Monographs 78(1):19–40.

(31) Sheldon RW, Prakash A, Sutcliffe Jr. WH (1972) The Size Distribution of Particles in the Ocean1. Limnology and Oceanography 17(3):327–340.

(32) Rall BC, et al. (2012) Universal temperature and body-mass scaling of feeding rates. Philosophical Transactions of the Royal Society B: Biological Sciences 367(1605):2923–2934.

(33) Saavedra S, Arroyo JI, Marquet PA, Kempes CP (2023) Linking metabolic scaling and coexistence theories.

(34) Schmiedel U, Oldeland J (2018) Vegetation responses to seasonal weather conditions and decreasing grazing pressure in the arid Succulent Karoo of South Africa. African Journal of Range and Forage Science 35(3-4):303–310.

(35) Creed RP (2006) Predator transitions in stream communities: a model and evidence from field studies. Journal of the North American Benthological Society 25(3):533–544.

(36) Landmann S, Holmes CM, Tikhonov M (2021) A simple regulatory architecture allows learning the statistical structure of a changing environment. eLife 10:e67455.

(37) Bruijning M, Metcalf CJE, Jongejans E, Ayroles JF (2020) The Evolution of Variance Control. Trends in Ecology & Evolution 35(1):22–33.

(38) De Bie T, et al. (2012) Body size and dispersal mode as key traits determining metacommunity structure of aquatic organisms. Ecology Letters 15(7):740–747.

(39) Woodward G, Warren P (2007) Body size and predatory interactions in freshwaters: scaling from individuals to communities in Body Size: The Structure and Function of Aquatic Ecosystems, Ecological Reviews, eds. Hildrew AG, Raffaelli DG, Edmonds-Brown R. (Cambridge University Press, Cambridge), pp. 98–117.

(40) Wilkes MA, et al. (2020) Trait-based ecology at large scales: Assessing functional trait correlations, phylogenetic constraints and spatial variability using open data. Global Change Biology 26(12):7255–7267.

(41) Tumolo BB, Albertson LK, Daniels MD, Cross WF, Sklar LL (2023) Facilitation strength across environmental and beneficiary trait gradients in stream communities. Journal of Animal Ecology 92(10):2005–2015.

(42) Wallace JB, Webster JR (1996) The role of macroinvertebrates in stream ecosystem function. Annual Review of Entomology 41:115–139.

(43) Smith DG, McBride GB (1990) New Zealand’s National Water Quality Monitoring Network - Design and First Year’s Operation1. JAWRA Journal of the American Water Resources Association 26(5):767–775.

(44) Quinn JM, Hickey CW (1990) Characterisation and classification of benthic invertebrate communities in 88 New Zealand rivers in relation to environmental factors. New Zealand Journal of Marine and Freshwater Research 24(3):387–409.

(45) Dolédec S, Phillips N, Townsend C (2011) Invertebrate community responses to land use at a broad spatial scale: trait and taxonomic measures compared in New Zealand rivers. Freshwater Biology 56(8):1670–1688.

(46) Chevene F, Doléadec S, Chessel D (1994) A fuzzy coding approach for the analysis of long-term ecological data. Freshwater Biology 31(3):295–309.

(47) Assaad CK, Devijver E, Gaussier E (2022) Survey and Evaluation of Causal Discovery Methods for Time Series. Journal of Artificial Intelligence Research 73:767–819.

(48) Günther W, Ninad U, Runge J (2023) Causal Discovery for time series from multiple datasets with latent contexts in Proceedings of the Thirty-Ninth Conference on Uncertainty in Artificial Intelligence. (PMLR), pp. 766–776.

(49) Kalisch M, Mächler M, Colombo D, Maathuis MH, Bühlmann P (2012) Causal Inference Using Graphical Models with the R Package pcalg. Journal of Statistical Software 47(11):1 –26. Section: Articles.

(50) Hauser A, Bühlmann P (2012) Characterization and greedy learning of interventional Markov equivalence classes of directed acyclic graphs. The Journal of Machine Learning Research 13(1):2409–2464.

(51) Tsamardinos I, Brown LE, Aliferis CF (2006) The max-min hill-climbing Bayesian network structure learning algorithm. Machine Learning 65(1):31–78.

(52) Kalisch M, Bühlmann P (2007) Estimating High-Dimensional Directed Acyclic Graphs with the PC-Algorithm. The Journal of Machine Learning Research 8:613–636.

(53) Palamara GM, Delius GW, Smith MJ, Petchey OL (2013) Predation effects on mean time to extinction under demographic stochasticity. Journal of Theoretical Biology 334:61–70.

(54) Gillespie DT (1977) Exact stochastic simulation of coupled chemical reactions. The Journal of Physical Chemistry 81(25):2340–2361.

